# MICAL1 activation by PAK1 mediates actin filament disassembly

**DOI:** 10.1101/2021.09.15.460490

**Authors:** David J. McGarry, Giovanni Castino, Sergio Lilla, Sara Zanivan, Michael F. Olson

## Abstract

The MICAL1 monooxygenase has emerged as an important regulator of filamentous actin (F-actin) structures that contribute to numerous processes including nervous system development, cell morphology, motility, viability and cytokinesis [1–4]. Activating *MICAL1* mutations have been linked with autosomal-dominant lateral temporal epilepsy, a genetic syndrome characterized by focal seizures with auditory symptoms [5], emphasizing the need for tight control of MICAL1 activity. F-actin binding to MICAL1 stimulates catalytic activity, resulting in the oxidation of actin methionine residues that promote F-actin disassembly [6, 7]. Although MICAL1 has been shown to be regulated via interactions of the autoinhibitory carboxyl-terminal coiled-coil region [8] with RAB8, RAB10 and RAB35 GTPases [9–12], or Plexin transmembrane receptors [13, 14], a mechanistic link between the RHO GTPase signaling pathways that control actin cytoskeleton dynamics and the regulation of MICAL1 activity had not been established. Here we show that the CDC42 GTPase effector PAK1 serine/threonine kinase associates with and phosphorylates MICAL1 on serine 817 (Ser817) and 960 (Ser960) residues, leading to accelerated F-actin disassembly. Deletion analysis mapped PAK1 binding to the amino-terminal catalytic monooxygenase and calponin domains, distinct from the carboxyl-terminal proteinprotein interaction domain. Stimulation of cells with extracellular ligands including basic fibroblast growth factor (FGF2) led to significant PAK-dependent Ser960 phosphorylation, thus linking extracellular signals to MICAL1 phosphorylation. Moreover, mass spectrometry analysis revealed that co-expression of MICAL1 with CDC42 and active PAK1 resulted in hundreds of proteins increasing their association with MICAL1, including the previously described MICAL1-interacting protein RAB10 [15]. These results provide the first insight into a redox-mediated actin disassembly pathway linking extracellular signals to cytoskeleton regulation via a RHO GTPase family member, and reveal a novel means of communication between RHO and RAB GTPase signaling pathways.

## Results

### MICAL1 binds active PAK1

High throughput phosphoproteomics studies have identified phosphorylated serine/threonine residues in MICAL1 [16–19]. Given the importance of serine/threonine kinases in the regulation of actin cytoskeleton organization by RHO GTPases [20–22], we sought to determine if MICAL1 was a target of RHO GTPase effector kinases. FLAG-tagged MICAL1 was expressed in HEK293T cells alone, coexpressed with myc-tagged constitutively-active CDC42 G12V or RHOA Q63L alone, or along with myc-tagged versions of the CDC42 effectors PAK1 or MRCKα, or the RHOA effectors ROCK1 or ROCK2 (**Figure 1A,** left panel). MICAL1 coimmunoprecipitated only with PAK1 in the presence of its activator CDC42 (**Figure 1A**, right panel). Western blotting PAK1 with a phospho-sensitive antibody against the activation loop Threonine 423 (T423) that must be phosphorylated for full activity [23] revealed that only CDC42-activated PAK1 associated with MICAL1 (**Figure 1B**). Treatment with increasing concentrations of the ATP-competitive Group I PAK inhibitor FRAX1036 [24] resulted in progressively reduced PAK1-MICAL1 association (**Figure 1C**). Furthermore, point mutations that blocked activation loop phosphorylation (T423A), reduced catalytic activity (K299R), or impaired CDC42 binding to the Cdc42/Rac interactive binding motif (CRIB) (H83L, H86L) blocked PAK1 activation by CDC42 and significantly reduced MICAL1 binding (**Figures 1D-E**). Since Group I PAK kinases undergo sequential conformational changes to become fully active, including CDC42-binding and activation loop phosphorylation [22], these results indicate that MICAL1 preferentially binds to PAK1 in its fully active conformation.

**Figure 1.**
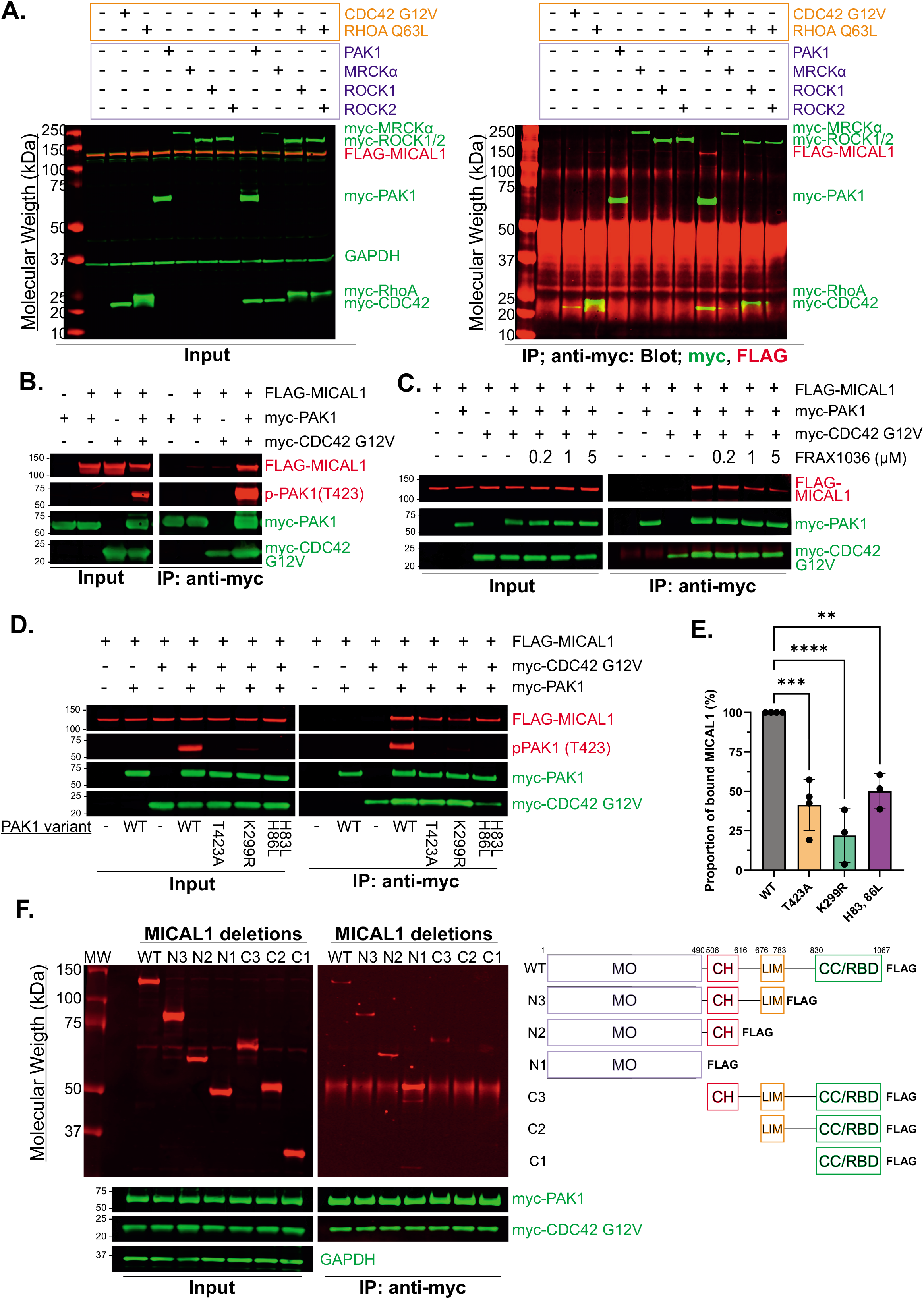
MICAL1 interacts with active PAK1. **A.** *Left panel*. Western blots of HEK293T cell lysates expressing FLAG-MICAL1 (red) in combination with myc-tagged CDC42 G12V or RHOA Q63L, without or with myc-tagged kinases PAK1, MRCKα, MRCKβ, ROCK1 or ROCK2 as indicated (green). GAPDH (green) served as a loading control. *Right panel*. Myc-tagged proteins were immunoprecipitated with anti-myc beads, then blotted for myc-tagged proteins with anti-myc antibody (green) and FLAG-MICAL1 with anti-FLAG antibody (red). **B.** *Left panel*. Western blot of HEK293T cell lysates expressing FLAG-MICAL1 without or with in combinations myc-PAK1 and myc-CDC42 G12V. *Right panel*. Myc-tagged proteins were immunoprecipitated with anti-myc beads, then blotted for myc-tagged proteins with anti-myc antibody (green), FLAG-MICAL1 with anti-FLAG antibody (red) and anti-PAK1 phospho-Threonine 423 (p-PAK1 (T423), red). **C.** *Left panel*. HEK293T cells expressing FLAG-MICAL1, myc-CDC42 G12V and myc-PAK1 were incubated for 18 hours with the PAK inhibitor FRAX1036 at the concentrations shown. *Right panel*. Myc-tagged proteins were immunoprecipitated with anti-myc beads, then blotted for myc-tagged proteins with anti-myc antibody (green) and FLAG-MICAL1 with anti-FLAG antibody (red). **D.** *Left panel*. HEK293T cells expressing myc-tagged PAK1 mutants without or with myc-tagged PAK1 and CDC42 G12V as indicated. *Right panel*. Myc-tagged proteins were immunoprecipitated from cells with anti-myc beads, then quantitatively Western blotted for myc-tagged proteins with anti-myc antibody (green) and FLAG-MICAL1 with anti-FLAG antibody by infrared scanning. **E.** Quantification of FLAG-MICAL1 co-immunoprecipitated with PAK1 mutants. Means ± SD, N = 3 independent experiments. ** = p<0.01, *** = p < 0.005, **** = p < 0.0001 (one-way ANOVA with *post-hoc* Dunnett’s multiple comparisons test). **F.** *Left panel*. Expression of FLAG-MICAL1 domain deletion mutants with myc-tagged PAK1 and CDC42 G12V. *Middle panel*. Immunoprecipitation of myc-tagged proteins and blotting for myc-tagged proteins with anti-myc antibody (green) and FLAG-MICAL1 with anti-FLAG antibody (red). *Right panel*. Schematic diagram of MICAL1 domains and the deletion constructs shown in the Western blot. MO; monooxygenase domain, CH; calponin homology domain, LIM; LIM domain, CC/RBD; coiled-coiled / RAB-binding domain.

MICAL1 is a multidomain protein, including catalytic monooxygenase (MO), calponin homology (CH), cysteine-rich Zinc-finger protein-protein interaction LIM, and coiled-coil RAB-binding (CC/RBD) domains (**Figure 1F**) [25, 26]. To identify PAK1-interacting regions, MICAL1 deletions were constructed that lacked one or more functional domain. When co-expressed with CDC42 G12V and PAK1, only the N1, N2, N3 and C3 constructs that all contained the MO and/or CH domains were co-immunoprecipitated (**Figure 1F**). Thus, unlike RAB GTPases or Plexin that bind to the carboxyl-terminal domain [10, 13, 14], PAK1 binds to the same MO + CH domains as F-actin [8, 27].

### PAK1 accelerates MICAL1 mediated F-actin disassembly

MICAL1 catalyzes redox-mediated F-actin depolymerization through the oxidation of methionine residues including Met44 and Met47 [6, 28]. Given recent structural evidence suggesting that MICAL1 exists in an autoinhibited state that can be relieved through protein-protein interactions [8, 29, 30], we hypothesized that PAK1 might regulate MICAL1 activity. To examine the effect of PAK1 on MICAL1 mediated F-actin disassembly, F-actin was incubated with either recombinant active PAK1 kinase domain (8.7 nM) or MICAL1 alone (133 nM), or in combination, for 5 minutes before ultracentrifugation separated the globular actin (G-actin) supernatant (S) and F-actin pelleted (P) fractions. Unpolymerized G-actin was entirely in the S fraction, while *in vitro* polymerization redistributed F-actin predominantly to the P fraction (**Figure 2A**). While the addition of active PAK1 alone caused no change in F-actin distribution, incubation with MICAL1 shifted some actin from P to S fractions (**Figure 2A**). The addition of active PAK1 to MICAL1 caused even greater shift of actin to the S fraction, indicating that PAK1 increased MICAL1 mediated F-actin disassembly. Changes in the fluorescence of pyrene-labelled actin was used to assess F-actin disassembly in real-time [31]. PAK1 alone did not affect F-actin levels over 25 minutes, while MICAL1 induced significant F-actin disassembly (**Figures 2B-C**). The addition of active PAK1 (4.35 nM) to MICAL1 (66.7 nM) resulted in significantly greater F-actin disassembly than was observed for MICAL1 alone (**Figure 2B-C**), in agreement with the observations from the fractionation study (**Figure 2A**). Taken together, these data indicate that MICAL1 activity is increased through its association with active PAK1.

**Figure 2.**
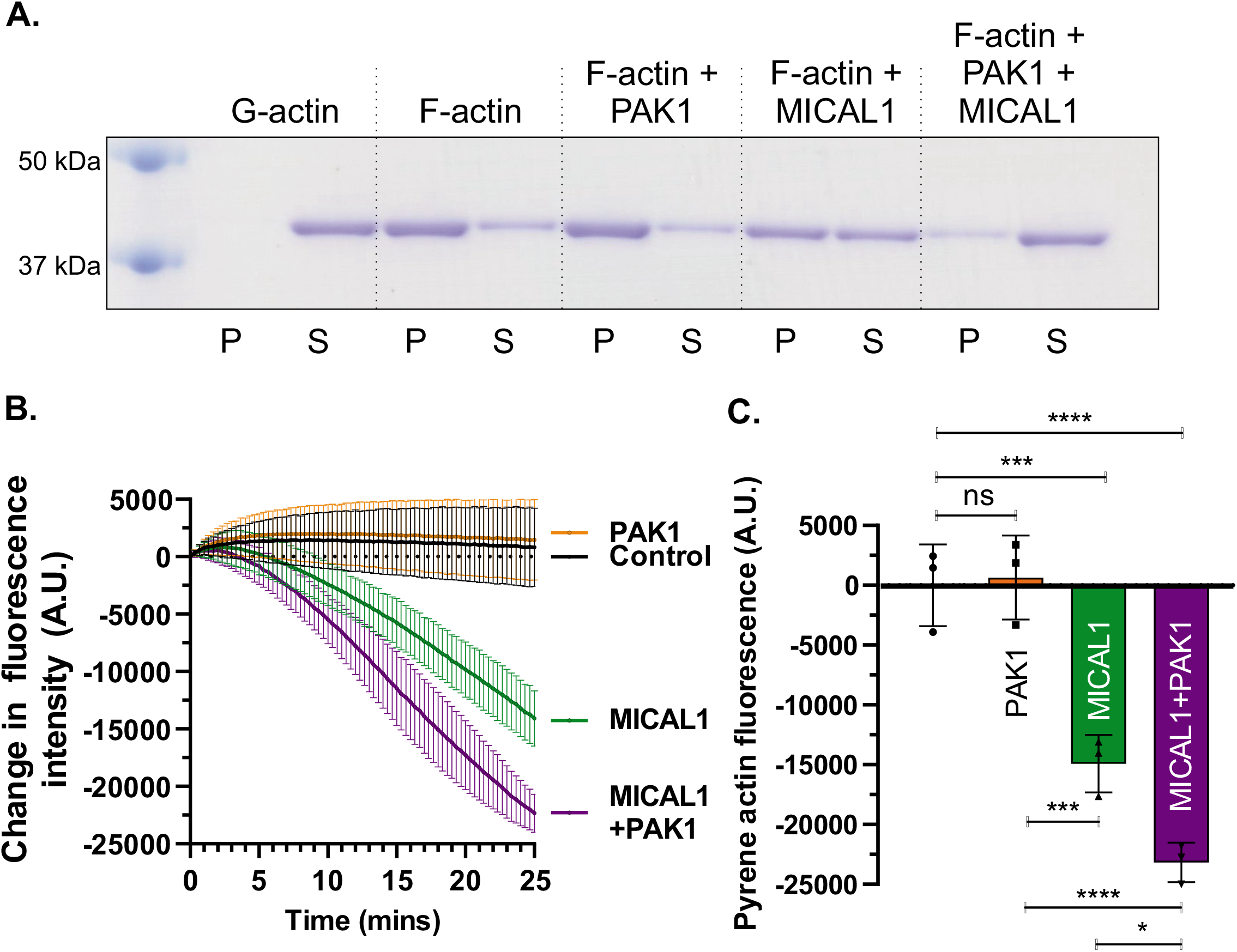
Active PAK1 promotes MICAL1 mediated F-actin disassembly. **A.** Actin sedimentation assay showing that active PAK1 increases MICAL1 mediated F-actin disassembly, indicated by the greater shift of F-actin from P (pellet) to S (supernatant) fractions. **B.** Pyrene-labelled F-actin depolymerization assay. Kinetic data showing MICAL1 mediated F-actin disassembly. Data from N = 3 independent experiments, means ± SD. **C.** Pyrene-labelled F-actin disassembly endpoint values at t = 25 mintues. Data show N = 3 independent experiments, means ± SD. * = p<0.05, ** = p<0.01, *** = p < 0.005, **** = p < 0.0001 (one-way ANOVA with *post-hoc* Dunnett’s multiple comparisons test).

### PAK1 phosphorylates MICAL1 at S817 and S960

PAK1 is a serine/threonine kinase that regulates cytoskeleton dynamics through the phosphorylation of downstream targets [32]. Given that MICAL1 binds to active PAK1 in a CDC42-dependent manner (**Figure 1**) and increases MICAL mediated Factin disassembly (**Figure 2**), we examined whether MICAL1 was a PAK1 substrate. Phosphorylation of FLAG-MICAL1 immunoprecipitated from HEK293T cells was detected with a pan-phospho-Ser/Thr antibody only after incubation *in vitro* with active recombinant PAK1 and 100 μM ATP (**Figure 3A**). While co-expression of FLAG-MICAL1 with wild-type PAK1 (PAK1^WT^) in HEK293T cells increased anti-phospho-Ser/Thr antibody staining of FLAG-MICAL1, catalytically inactive PAK1 K299R (PAK1^K299R^) did not (**Figure 3B**). Additionally, treatment of HEK293T cells with Group I PAK inhibitor FRAX1036 reduced FLAG-MICAL1 phosphorylation by CDC42 activated PAK^WT^ in a dose dependent manner (**Figure 3C**). To identify sites of MICAL1 phosphorylation, FLAG-MICAL1 was immunoprecipitated from cells without or with co-expressed CDC42 G12V plus PAK1^WT^, followed by tryptic digestion and tandem mass spectrometry (MS/MS). Phosphoproteomic analysis identified serine 817 (S817) and serine 960 (S960) as the only sites of 10 serine or threonine residues identified that were consistently increased in their phosphorylation in cells co-expressing MICAL1 with active CDC42 and PAK1 (**Figure 3D, Figure S1A-B**). S817 is located near the end of an α-helix composed of residues 809-818 (**Figure S1B**, inset) [33] between the LIM and CC/RBD domains (**Figure 3E**). S960 is at the end of the first of the three α-helices that comprise the carboxyl-terminal coiled-coiled/RBD domain (**Figure 3E**) [15]. Although neither site is conserved in the closely-related human MICAL2 or MICAL3 proteins, they are both highly conserved across mammalian species (S817 = 31/31, S960 = 28/31 example mammals; **Supplemental Table 1**).

**Figure 3.**
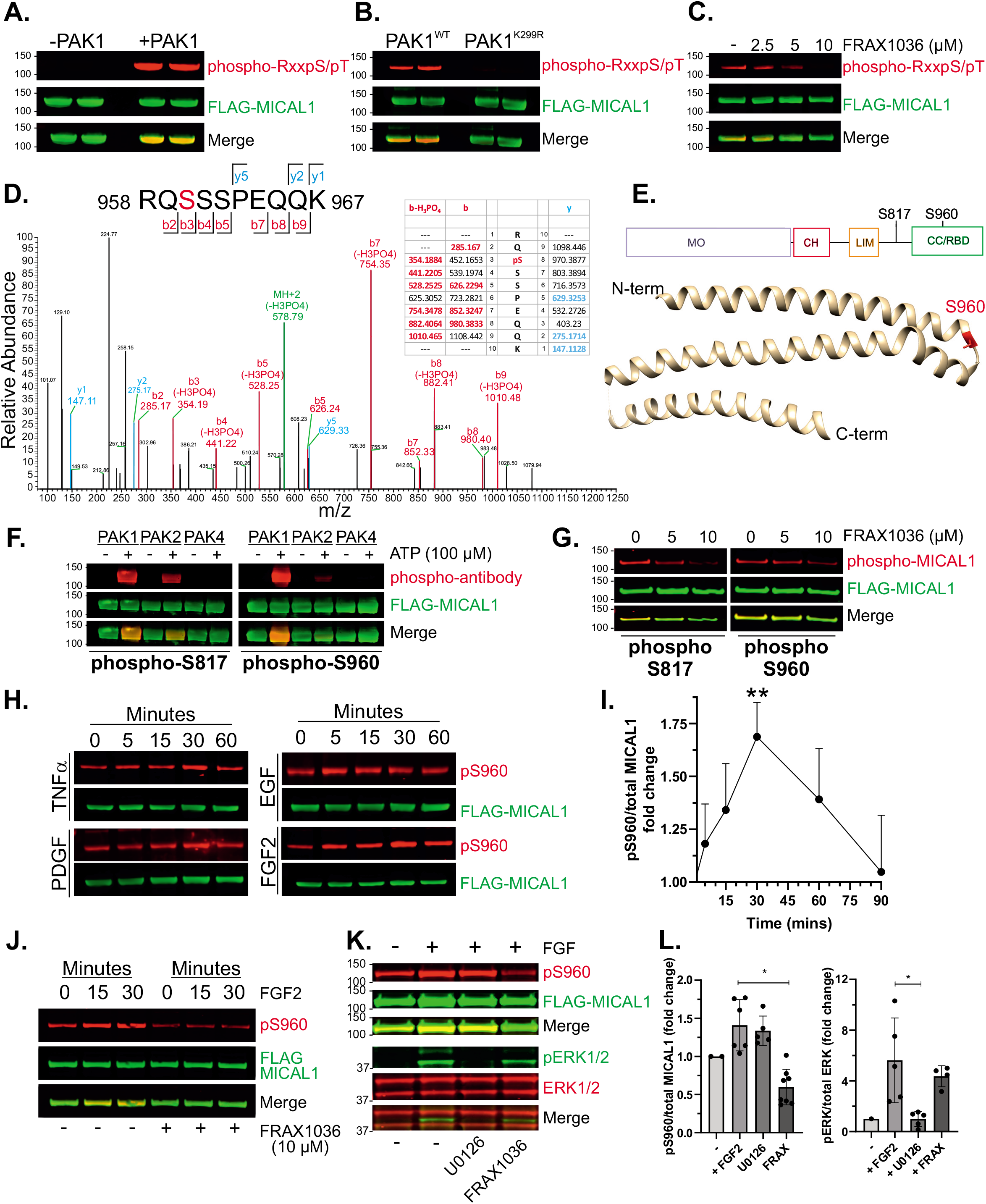
Ser817 and Ser960 MICAL1 are PAK1 substrates. **A.** *In vitro* phosphorylation of FLAG-MICAL1 by recombinant active PAK1. FLAG-MICAL1 was immunoprecipitated from transfected HEK293T cells with anti-FLAG beads. Proteins were resuspended in kinase buffer and incubated with or without 50 ng PAK1 for 1 hr at 30°C. Samples were boiled in 1x LDS reducing sample buffer and proteins resolved by Western blotting with anti-phospho-RxxpS/pT antibody (red) or anti-FLAG antibody (green). **B.** Immunoprecipitation of FLAG-MICAL1 co-expressed with CDC42 G12V and either wild-type (PAK1^WT^) or kinase-dead PAK1 (PAK1^KD^) from transfected HEK293T cells. Samples were boiled in 1x LDS reducing sample buffer and proteins resolved by Western blotting with anti-phospho-RxxpS/pT antibody (red) or anti-FLAG antibody (green). **C.** HEK293T cells were transfected with FLAG-MICAL1, CDC42 G12V and myc-PAK1^WT^ before overnight incubation with the PAK1 inhibitor FRAX1036. FLAG-MICAL1 was immunoprecipitated with anti-FLAG beads and proteins resolved by Western blotting with anti-phospho-RxxpS/pT antibody (red) or anti-FLAG antibody (green). **D.** Higher energy collision dissociation (HCD) MS/MS fragmentation spectra of MICAL1 tryptic peptide 958-967 containing S960 (red). The b and y ion series ions identified experimentally are indicated in blue and red, respectively. **E.** Schematic diagram of MICAL1 domains: monooxygenase (MO), calponin homology (CH) LIM, coiled-coil Rab-binding domain (CC/RBD), showing location of S817 and S960 phosphorylation sites. UCSF Chimera [51] modeling of the MICAL1 C-terminal coiled-coil domain indicates the S960 (red) location (Protein data base reference 5LE0) [15]. **F.** *In vitro* FLAG-MICAL1 phosphorylation by recombinant PAK1, PAK2 and PAK4. FLAG-MICAL1 was immunoprecipitated from transfected HEK293T cells and incubated with 50 ng of active PAK1, PAK2 or PAK4. Assays were performed with or without ATP (100 μM) to assess the specificity of phosphorylation. Western blotting was with antibodies against FLAG-MICAL1 (green) and pS817 (left panels; red) or pS960 (right panels; red). **G.** HEK293T cells expressing FLAG-MICAL1, myc-CDC42 G12V and myc-PAK1 were incubated for 18 hours with the PAK inhibitor FRAX1036 at the indicated concentrations. Proteins were resolved by Western blotting with anti-phospho-S817 (left panels, red) or anti-phospho-S960 (right panels, red) antibody, and anti-FLAG antibody (green). **H.** HEK293T expressing FLAG-MICAL1 were serum starved overnight before stimulation with 20 ng/ml TNFα, EGF, PDGF or FGF2 for the indicated times. Proteins were resolved by Western blotting with anti-phospho-S960 antibody (red) and anti-FLAG antibody (green). **I.** Quantification of FGF2 stimulated S960 phosphorylation. Data show means ± SD, N = 3 independent experiments. ** = p<0.01 (one-way ANOVA with *post-hoc* Dunnett’s multiple comparisons test). **J.** HEK293T expressing FLAG-MICAL1 cells were stimulated with FGF2 without or with 10 μM FRAX1036 for the indicated times. Proteins were resolved by Western blotting with anti-phospho-S960 antibody (red) and anti-FLAG antibody (green). **K.** HEK293T expressing FLAG-MICAL1 cells were stimulated with FGF2 for 30 minutes with or without 10 μM U012MEK inhibitor or 10 μM FRAX1036. Proteins were resolved by Western blotting with anti-phospho-S960 (red) and anti-FLAG antibodies (green), or with anti-phospho-Thr202/Tyr204 ERK1/2 (green) and total ERK1/2 antibodies (red). **L.** Quantification of FGF2 stimulated S960 phosphorylation with MEK or PAK inhibition. Data show means ± SD, N = 4-8 independent experiments. * = p<0.05 (one-way ANOVA with *post-hoc* Dunnett’s multiple comparisons test).

The phospho-specificity of polyclonal antibodies raised against phosphorylated S817 (pS817) and S960 (pS960) sites were confirmed by dot blotting phosphorylated (P) or unphosphorylated (U) S817 or S960 peptides (**Figure S1C**). Western blotting FLAG-MICAL1 serine to alanine mutants (S817A, S960A) coexpressed with wild-type (WT) or kinase-dead (KD) PAK1 in HEK293T cells established the phospho-specificity of these antibodies and lack of cross reactivity (**Figure S1D**). Incubation of immunoprecipitated FLAG-MICAL1 with group I PAK1 or PAK2, or group II PAK4 recombinant kinases in the absence or presence of 100 μM ATP revealed that only group I PAK1 and PAK2 kinases phosphorylated S817 and S960 (**Figure 3F**). Furthermore, S817 and S960 phosphorylation by endogenous PAK were inhibited by the group I PAK inhibitor FRAX1036 (**Figure 3G**). To determine if S817 or S960 phosphorylation altered MICAL1 subcellular localization, proximity ligation assays (PLA) were performed in transfected U2OS human bone osteosarcoma epithelial cells (**Figure S2A**). Total MICAL1 expression was detected using pairs of mouse and rabbit anti-FLAG antibodies, which was equivalent when co-expressed with PAK^WT^ or PAK^KD^ (**Figure S2B**). PLA signals for rabbit pS817 or pS960 antibodies paired with mouse anti-FLAG antibody were only detected in PAK^WT^ expressing cells, consistent with their phospho-sensitivity (**Figure S2B**). High resolution confocal imaging did not reveal noticeable differences in the distribution of total MICAL1 co-expressed with PAK^KD^ compared to MICAL1 S817 or S960 phosphorylated by PAK^WT^ from their predominantly cytoplasmic locations, suggesting that PAK-mediated phosphorylation does not have a large effect on MICAL1 subcellular localization (**Figure S3**).

PAK1 is activated by extracellular stimuli [22, 34], suggesting that MICAL1 phosphorylation might be similarly regulated. Serum-starved HEK293T cells expressing FLAG-MICAL1 were stimulated for varying times up to 60 minutes with tumour necrosis factor α (TNFα), epidermal growth factor (EGF), platelet-derived growth factor (PDGF) or basic fibroblast growth factor (FGF2). Western blotting revealed increasing pS960 staining that peaked 30 minutes after TNFα, PDGF or FGF2 treatment, and after 5 minutes with EGF stimulation (**Figure 3H**). FGF2 significantly increased S960 phosphorylation by ~60% after 30 minutes, with a return to unstimulated levels after 90 minutes (**Figure 3I**). The increase in FGF2-stimulated S960 phosphorylated was effectively blocked by the group I PAK inhibitor FRAX1036 (**Figure 3J**). To confirm that the inhibition of S960 phosphorylation by FRAX1036 occurred downstream of the FGF2 receptor, the effects of FRAX1036 and the mitogen-activated protein kinase kinase (MEK) inhibitor U0126 [35] were each evaluated on MICAL1 phosphorylation and mitogen-activated protein kinase (MAPK) pathway activation. While U0126 had no effect on FGF2-stimulated MICAL1 S960 phosphorylation and significantly inhibited ERK1 and ERK2 Thr202/Tyr204 phosphorylation to unstimulated levels, FRAX1036 significantly reduced MICAL1 S960 phosphorylation levels to below those observed in unstimulated cells but did not affect ERK1 and ERK phosphorylation (**Figures 3K-L**). The lack of effect of FRAX1036 on FGF2-stimulated MAPK activation indicates that the reduction in MICAL1 S960 phosphorylation occurred downstream of the receptor for FGF2 at the level of PAK activity. Taken together, these results demonstrate that stimulation with extracellular peptide ligands leads to MICAL1 phosphorylation.

### MICAL1 phosphorylation by PAK1 promotes RAB10 binding

Structural analysis shows that the majority of MICAL1 exists in a closed inactive conformation autoinhibited by the carboxyl-terminal region, which can be relieved by the binding of proteins such as RAB GTPases to the CC/RBD region to open and activate MICAL1 [8, 10]. The affinity of the CC/RBD for RAB proteins was found to be dependent on the positions of the first and second α-helices [10], raising the possibility that PAK1 phosphorylation of S817, located between the LIM and CC/RBD protein binding domains, and S960, located at the end of α-helix1 that leads to a turn that couples to the start of α-helix2 (**Figure 3E**), might affect protein binding. To identify MICAL1 interacting proteins affected by PAK1 phosphorylation, FLAG-MICAL1 co-expressed with active CDC42 and PAK1^KD^ or PAK1^WT^ was immunoprecipitated, followed by tryptic digestion, quantification and MS/MS analysis of co-precipitated proteins. The volcano plot in **Figure 4A** depicts the greater number of proteins associated with MICAL1 expressed with PAK1^WT^ versus PAK^KD^ (216 decreased vs 473 increased). The magnitudes of the difference in individual proteins binding to phosphorylated MICAL1 was also significantly greater for increased proteins (median = 0.53, mean = 0.76) than decreased proteins (median = 0.32, mean = 0.48) (**Figure 4B**). MS/MS analysis revealed increased association of both PAK1 and CDC42 (**Figure 4A** and **Supplementary Table 2**), in agreement with results in **Figure 1**. In addition, the previously identified MICAL1 interacting proteins RAB7A and RAB10 were also identified as being increased in their binding to phosphorylated MICAL1 (**Figure 4A**). Interestingly, the phosphoserine-binding protein YWHAE (14-3-3ε) was also bound to a greater degree to MICAL1 co-expressed with CDC42 and PAK1^WT^ (**Figure 4A**), consistent with PAK1 increasing the phosphorylation of S817 and S960. The cluster diagram in **Figure 4C** depicts the most significantly enriched interactors that were increased in their binding to phosphorylated MICAL1, with the proximity to the center being indicative of the statistical significance of the interaction.

**Figure 4.**
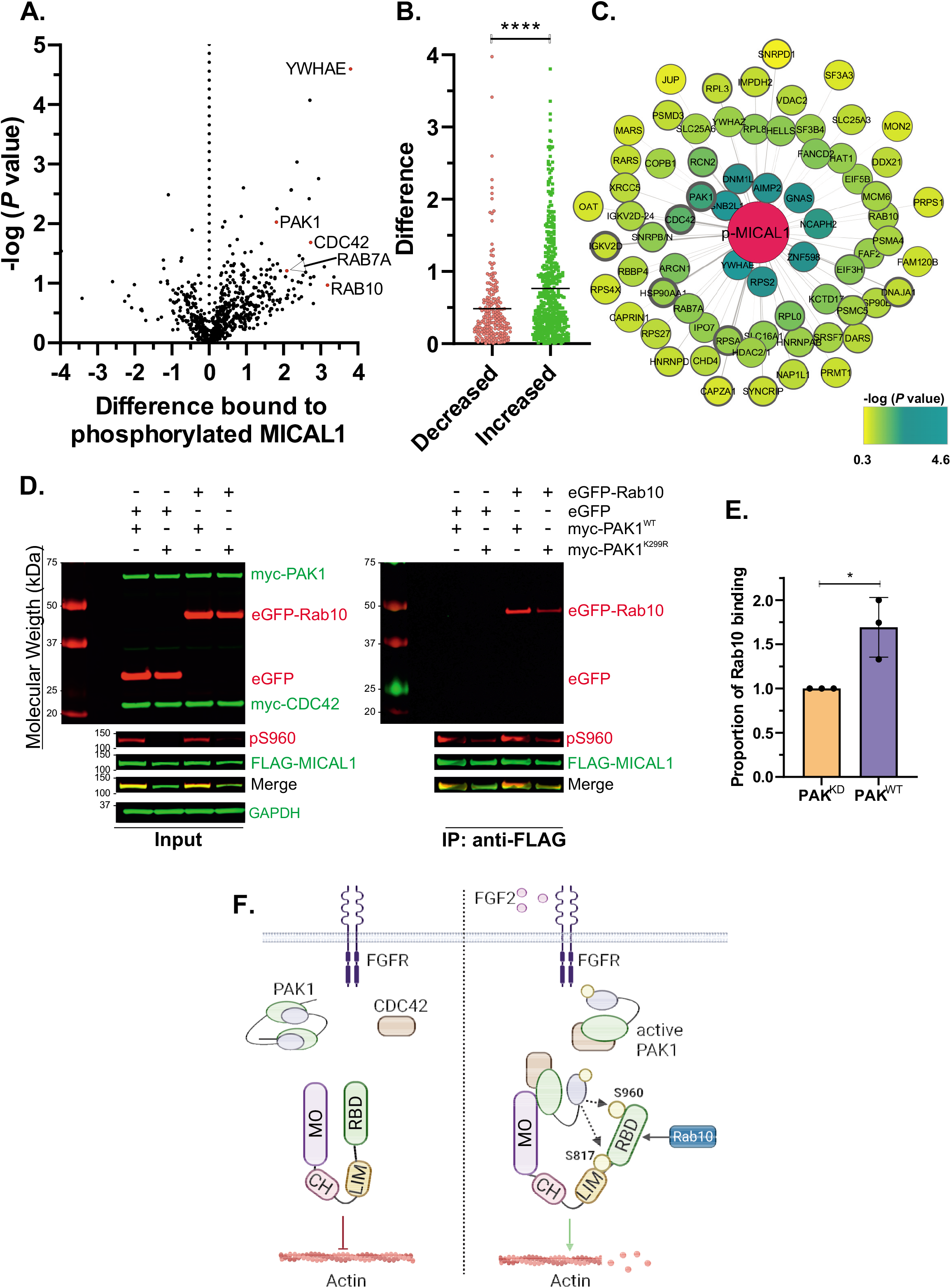
Phosphorylation of MICAL1 recruits RAB10a binding. **A.** Volcano plot showing proteins that were increased (positive values) in their association with phosphorylated MICAL1 expressed with CDC42 G12V and PAK1^WT^ relative to their association with MICAL1 expressed with CDC42 G12V and PAK1^KD^ (negative values). Proteins described in the manuscript are indicated with red dots. **B.** The differences in proteins preferentially associated with phosphorylated MICAL1 expressed with CDC42 G12V and PAK1^WT^ (increased; green dots) relative to their association with MICAL1 expressed with CDC42 G12V and PAK1^KD^ (decreased; red dots). Means indicated by the solid line. **** = p<0.0001 (Student’s t-test; unpaired, two-tailed). **C.** Cytoscape [49] network shows the interactions of individual proteins with phosphorylated MICAL1, where distance to the center is indicative of the statistical significance of the interaction, and the thickness of the circle boundaries is related to the Protein Andromeda scores taken from MaxQuant. **D.** *Left panel*. HEK293T cells were transfected with FLAG-MICAL1, myc-CDC42 G12V, eGFP, eGFP-RAB10a and myc-tagged PAK1^WT^ or PAK1^KD^. Proteins were resolved by Western blotting with anti-myc tag antibody (green), anti-eGFP-tag antibody (red), FLAG-MICAL1 with anti-FLAG antibody (green), anti-phospho-S960 MICAL1 antibody (red) and anti-GAPDH antibody (green). *Right panel*, FLAG-MICAL1 was immunoprecipitated using anti-FLAG beads, then blotted for eGFP-tagged proteins (red), FLAG-MICAL1 with anti-FLAG antibody (green) and anti-phospho-S960 antibody (red) **E.** Quantification of RAB10 binding to MICAL1 expressed with myc-CDC42 G12V and myc-tagged PAK1^WT^ or PAK1^KD^. Data show means ± SD., N = 3 independent experiments. * = p<0.05 (Student’s t-test; unpaired, two-tailed). **F.** *Left side*; In quiescent cells, MICAL1 exists in an autoinhibited state through interactions between N- and C-termini [8]. Inactive PAK1 is a homodimer [52, 53] in the absence of activated CDC42. *Right side*; In response to FGF2 stimulation, CDC42 activates PAK1 leading to PAK1 binding to the MICAL1 monooxygenase (MO) and calponin homology (CH) domains. PAK1 subsequently phosphorylates MICAL1 at S817 and S960. These phosphorylations result in: 1) increased RAB10 binding to the C-terminal Rab-binding domain (RBD) and 2) increased MICAL1-mediated F-actin disassembly.

Of particular interest was the identification of RAB10 as being preferentially associated with phosphorylated MICAL1 (**Figures 4A** and **4C**). Although several lines of evidence have shown that MICAL1 associates with RAB GTPases *in vitro* [10], in cells [12] and in yeast two-hybrid screens [9], there has been no indication that MICAL1 post-translational modifications influence RAB GTPase binding. To determine whether MICAL1 phosphorylation affected RAB10 binding, FLAG-MICAL1 co-expressed with CDC42 and PAK1^WT^ or PAK1^KD^ was also expressed with enhanced green fluorescent protein (eGFP) or eGFP-RAB10 (**Figure 4D**). MICAL1 was phosphorylated on S960 when co-expressed with CDC42 and PAK1^WT^ (**Figure 4D**, left panel). Immunoprecipitation of FLAG-MICAL1 resulted in the specific coimmunoprecipitation of eGFP-RAB10, which was significantly greater when S960 was phosphorylated by PAK1 (**Figure 4D**, right panel; **Figure 4E**). These results reveal for the first time that there is a two-way relationship between MICAL1 and RAB GTPase binding; in addition to RAB GTPase binding inducing conformational changes that increase activity [10, 12], MICAL1 phosphorylation by PAK1 increases RAB GTPase binding.

## Discussion

For over 30 years, RHO GTPases have been situated in central roles in the regulation of actin cytoskeleton organization required for many cellular processes such as cell movement, polarity, adhesion, and membrane transport [36]. Despite decades of study, the full repertoire of mechanisms that RHO GTPases use to regulate cytoskeleton dynamics has not been established. MICAL1 has emerged as an important regulator of filamentous actin structures, with demonstrated roles downstream of Semaphorin/Plexin signalling [7, 14]. However, a role for MICAL1 in RHO GTPase signalling pathways had not been demonstrated. We now reveal that PAK1 activation by CDC42 results in the association with and phosphorylation of MICAL1 on S817 and S960, leading to increased F-actin disassembly (**Figure 4F**). Mapping the interacting MICAL1 domains revealed that PAK1 associates with the monooxygenase and calponin homology domains, similar to F-actin [8, 27]. Given that MICAL1 has been proposed to be autoinhibited by the carboxyl-terminal region [8, 10], the closed protein confirmation could bring S817 and S960 into proximity to PAK1 associated with the MO + CH domains (**Figure 4F**). PAK1 phosphorylation of MICAL1 would then be predicted to relax the closed conformation to enable increased F-actin depolymerization by the catalytic regions and increased RAB GTPase binding to the CC/RBD domain, which could further contribute to MICAL1 activation.

Although PAK1-mediated phosphorylation did not affect bulk PAK1 localization as determined by proximity ligation assays, the additional proteins binding to phosphorylated MICAL1 revealed by mass spectrometry analysis might act to concentrate activated MICAL1 at discrete sites to locally dissemble F-actin. Consistent with this possibility, previous studies have shown important roles for the targeting of MICAL proteins to specific cytoskeleton structures via interactions with myosin 5A [33], myosin 15 [37] and coronin 1C [38, 39]. The increased binding of RAB7A and RAB10 to phosphorylated MICAL1 suggests a role for PAK1 in enabling RAB-mediated membrane trafficking [40, 41], evidence of a novel link between RHO and RAB GTPase signalling pathways.

## Methods

### Cell culture

HEK293T cells were grown in DMEM (Gibco) supplemented with 10% fetal bovine serum (FBS, Gibco), 2 mM L-glutamine (Gibco), 10 U/ml penicillin and 10 μg/ml streptomycin (Gibco) and kept at 37°C with 5% CO2 in a humidified incubator. FreeStyle™ 293-F cells were grown in FreeStyle™ 293 Expression Medium (Gibco) under constant agitation at 37 °C with 5% CO2 in a humidified incubator.

### Plasmids

pcDNA3.1-MICAL1-FLAG was generated by amplifying MICAL1 from eGFP-MICAL1 (a kind gift from Prof. Steve Caplan, University of Nebraska Medical Center, Omaha, USA) together with a C-terminal FLAG tag, and cloning it into pcDNA3.1 (5’-actgaaggatccgccaccatggcttcacctacctccacc −3’, 5’-actgaagcggccgcttacttgtcgtcatcgtctttgtagtcgccctgggcccctgtccccaa −3’). Plasmids harbouring MICAL1 domain mutants were generated using Q5^®^ Site-Directed Mutagenesis Kits (New England BioLabs) according to the manufacturer’s protocol, with the following primer pairs; N1 (1-490 amino acids, 5’-catcgtctttgtagtcaggctccttggctagc-3’, 5’-gctagccaaggagcctgactacaaagacgatg-3’), N2 (1-616 amino acids, 5’-caagagcatggcccacgactacaaagacgatga-3’, 5’-tcatcgtctttgtagtcgtgggccatgctcttg-3’), N3 (1-783 amino acids, 5’-catcgtctttgtagtctgagaggcctggtggc-3’, 5’-gccaccaggcctctcagactacaaagacgatg-3’), C1 (830-1067 amino acids, 5’-ggaggcttgggtggcatggtggcggatc-3’, 5’-gatccgccaccatgccacccaagcctcc-3’), C2 (676-1067 amino acids, 5’-gatccgccaccatgccagagcctggtgt-3’, 5’-acaccaggctctggcatggtggcggatc-3’), and C3 (506-1067 amino acids, 5’-gcctgccgaccccatggtggcgga-3’, 5’-tccgccaccatggggtcggcaggc-3’). Plasmids for the expression of MICAL1 Ser to Ala mutants (S817A, S960A), were generated using Q5^®^ Site-Directed Mutagenesis Kits (New England BioLabs) according to the manufacturer’s protocol, with the following primer pairs; S817A (5’-ccagcggttggcctcccttaacc-3’, 5’-cgctccgggctggagagg-3’), S960A (5’-gaggcgccaggccagttccccag-3’, 5’-aaggccagctccagcttc-3’). The double S817A/S960 mutant was generated from the S817A mutant using the S960A primer pair mentioned above.

Plasmids for the expression of myc-PAK1 activity mutants (T423A, K299R, H83L/H86L), were generated from pCMV6M-myc-PAK1 using Q5^®^ Site-Directed Mutagenesis Kits (New England BioLabs) according to the manufacturer’s protocol, with the following primer pairs; T423A (5’-caaacggagcgccatggtaggaa-3’, 5’-ctctgctctggggttatctg-3’), K299R (5’-ggtggccattaggcagatgaatc-3’, 5’-tcctgtcctgtggccaca-3’) and H83L/H86L (5’-attctcgtcggttttgatgctgtc-3’, 5’-tgtgagttcaaaatctgaagggagag-3’).

### Protein expression and purification

Full-length human MICAL1 protein was expressed in FreeStyle™ 293-F cells. A 12xHis tag was cloned into the N-terminus of pcDNA3.1-MICAL1-FLAG using Q5^®^ Site-Directed Mutagenesis Kits (New England BioLabs) according to the manufacturer’s protocol, with the following primer pairs; 5’-ccaccacgagaatttgtattttcagggtgcttcacctacctccacc-3’ 5’-tgatgatgatggtggtggtgatgatgatgcatggtggcggatccgag-3’. Prior to transfection, cells were grown to a density of 1×10^6^ cells/ml. For a 0.5 litre culture, cells were transfected with 0.625 mg of pcDNA3.1-12xHis-MICAL1-FLAG using polyethylenimine (PEI) and left to express for 3 days.

For purification, cells were pelleted at 2000 rpm for 10 minutes and washed 2x in ice cold PBS. Cells were lysed by sonication in 20 ml of lysis buffer (50 mM Tris pH 8.0, 300 mM NaCl, 2 mM MgCl_2_, 20 mM imidazole, 1x Complete protease inhibitor (Roche)), before lysates centrifuged at 10,000 xG for 30 minutes. His-tagged MICAL1 was purified using HisTrap HP columns (GE Healthcare). Proteins were further purified using size exclusion chromatography and stored in a buffer of 50 mM Tris pH 8.0, 300 mM NaCl, 1 mM MgCl_2_, and 1 mM β-mercaptoethanol.

### Cell transfection, lysis and immunoprecipitation

HEK293T cells were plated in 100 mm dishes at 3×10^6^ cells per dish. The following day, media was replaced with OptiMEM and cells were transfected with 10 μg DNA using Lipofectamine (Invitrogen, 18324012), according to the manufacturer’s protocol. After 6 hours, the transfection media was replaced with cell culture media and cells were left to grow overnight. Lysis and immunoprecipitation of FLAG and myc-tagged constructs are as described previously [42]. Cells were washed with ice cold PBS and lysed in ice-cold TBS buffer (pH 7.5) containing 1 mM EDTA, 1% (v/v) Triton X-100, 20 mM NaF, 20 mM β-glycophosphate, 0.2 μM Na_3_VO_4_, 20 μg/mL aprotinin and 1x Complete protease inhibitor (Roche). Lysates were incubated on a rotating wheel for 30 minutes at 4°C, before centrifugation at 13,000 rpm for 10 minutes. Protein concentration was measured using the BCA assay and lysates were subsequently prepared for Western blot analysis or for immunoprecipitation reactions.

For immunoprecipitation reactions, lysates were incubated with anti-myc agarose beads (Sigma, A7470) or anti-FLAG agarose beads (Sigma, A2220) for 2 hours at 4°C. Beads were washed 3x in TBS buffer prior to downstream processing. For western blot analysis, beads were boiled in 1x LDS sample buffer (Invitrogen, B0007) containing 10 mM DTT for 5 minutes at 95°C, after which beads were centrifuged for 13,000 rpm for 1 min and supernatants collected.

### *In vitro* phosphorylation assays

*In vitro* phosphorylation assays were performed as previously described [42]. For assays involving FLAG-MICAL1 bound to anti-FLAG beads, beads were washed 1x in phosphatase buffer (P0753; New England BioLabs) for 5 minutes at 4°C. In order to remove prior phosphorylations, beads were pelleted and resuspended in 100 μl of phosphatase buffer with lambda protein phosphatase (P0753; New England Biolabs) before agitation at 30°C for 30 minutes. After phosphatase treatment, MICAL1-bound beads were washed 3x in PBS and 1x in kinase buffer (20 mM Tris pH 7.5, 0.5 mM MgCl_2_, 0.01% (v/v) Tween-20). For assays involving phosphorylation of recombinant MICAL1, MICAL1 protein was washed 3x in TBS using a 50 kDa cut off filter (Amicon, UFC505008) prior to lambda protein phosphatase treatment as outlined above. After phosphatase treatment, recombinant MICAL1 protein was washed 3x in PBS using a 50 kDa filter to remove any residual lambda protein phosphatase (~25 kDa).

For PAK1 phosphorylation assays, beads or recombinant protein (333 nM) were incubated in 150 μl kinase buffer supplemented with 100 μM ATP and 150 ng (21.7 nM) of active, recombinant PAK1 (Millipore, 14-927) and agitated at 30°C for 1 hr. To stop the reaction, samples were boiled in 1x LDS buffer for 5 minutes at 95°C.

### Proteolytic digestion of proteins ‘in gel’

The region containing MICAL1 was excised from gel and washed twice with 50 mM ammonium bicarbonate, and 50 mM ammonium bicarbonate with 50% acetonitrile. Proteins in the gel bands were then reduced using dithiothreitol (10 mM at 54°C for 30 minutes) and subsequently alkylated with iodoacetamide (55 mM at room temperature for 45 minutes). Gel pieces were washed again with 50 mM ammonium bicarbonate; 50 mM ammonium bicarbonate with 50% acetonitrile; and finally dehydrated using acetonitrile before drying in a SpeedVac. Trypsin (5 μg/ml trypsin gold (Promega) in 25 mM ammonium bicarbonate) was added and incubated for 12 h at 35 °C. Tryptic peptides were extracted from gel pieces with two washes of 50% (v/v) acetonitrile/water + 1% trifluoroacetic acid (TFA) and subsequently dried in a SpeedVac. Dried peptides were resuspended in 0.1% TFA and desalted using StageTips [43].

### MS analysis

Tryptic peptides were separated by nanoscale C18 reverse-phase liquid chromatography using an EASY-nLC 1200 (Thermo Fisher Scientific) coupled online to an Orbitrap Q-Exactive HF mass spectrometer (Thermo Fisher Scientific) via nanoelectrospray ion source (Sonation). Peptides were separated on a 50 cm fused silica emitter (New Objective) packed in house with reverse phase Reprosil Pur Basic 1.9 μm (Dr. Maisch GmbH). For the full scan, a resolution of 60,000 at 250 Th was used. The top fifteen most intense ions in the full MS were isolated for fragmentation with a target of 50,000 ions at a resolution of 15,000 at 250 Th. MS data were acquired using the XCalibur software (Thermo Fisher Scientific). An Active Background Ion Reduction Device (ABIRD) was used to decrease ambient contaminant signal level.

### Data analysis

MaxQuant software [44] version 1.5.5.1 was used to process MS Raw files and searched with Andromeda search engine [45], querying UniProt [46] Homo sapiens database (09/07/2016; 92,939 entries). Specificity for trypsin cleavage and maximum two missed cleavages were requested for the search. Cysteine carbamidomethylation was set as fixed modification. Methionine oxidation, Serine Threonine and Tyrosine Phosphorylation and N-terminal acetylation were specified as variable modifications. The peptide and protein false discovery rate (FDR) was set to 1%.

MaxQuant outputs were analysed with Perseus software [47] version 1.5.5.3. The MaxQuant output “Phospho (STY) Sites.txt” file was used for quantification of phosphorylated peptides. Reverse and Potential Contaminant peptides, as specified by MaxQuant, were removed. For phospho-proteomic studies, only phosphorylated peptides having: score difference greater than 5, a localisation probability higher than 0.75, and robust quantification in three out of three replicate experiments were included in the analysis. For protein enrichment studies, only protein groups identified with at least one uniquely assigned peptide were used for the quantification. For label-free quantification, proteins quantified in all five replicates in at least one group, were measured according to the label-free quantification algorithm available in MaxQuant [48]. Significantly enriched proteins were selected using a t-test with a 5% FDR (permutation based).

Volcano plots were generated with Perseus software [47] version 1.5.5.3. For cluster visualisation, proteins that showed a significant difference in binding to phosphorylated MICAL1 based on Volcano plot analysis were visualised using Cytoscape [49].

### Dot blots

For each phospho-S817 and phospho-S960 antibody, two peptides were synthesised by GenScript. One peptide contained a phosphorylated serine (P); 817 – RQRL(pS)SLNLTPDPEC, 960 – RRQ(pS)SSPEQQKKLWC, and the other contained a non-phosphorylated serine (U); 817 – RQRLSSLNLTPDPEC, 960 – RRQSSSPEQQKKLWC. A dose range of peptides were spotted onto a nitrocellulose membrane that had been pre-soaked in TBS + 0.01% Tween-20 (TBST) and dried. The membranes were blocked in 0.5% BSA/TBST for 30 min and incubated with the phospho-S817 or phospho-S960 antibodies (1:500) overnight at 4°C in 0.5% BSA/TBST. The following day, membranes were washed in TBST and incubated for 1 hr at room temperature with a secondary rabbit-HRP antibody (1:50,000). Membranes were developed using SuperSignal™ West Femto Maximum Sensitivity Substrate (Thermo Scientific, 34095).

### Western blots

Western blots were performed as described previously [50]. The following primary antibodies were used in the study; rabbit anti-FLAG (F7425, Sigma), mouse anti-FLAG (F1804, Sigma), mouse anti-myc (M4439, Sigma), rabbit anti-phospho-PAK1 T423 (2601S, Cell Signaling), mouse anti-GAPDH (MAB374, Millipore), rabbit anti-phospho-RxxS*/T* (9614S, Cell Signaling), mouse anti-phospho ERK1/2 (9106S, Cell Signaling), rabbit anti-ERK (9102S, Cell Signaling), rabbit anti-GFP (G1544, Sigma). The secondary antibodies used were IRDye^®^ 680RD and 800CW (LiCor). Proteins were visualized by infrared imaging using LiCor Odyssey CLx.

### Actin sedimentation

Actin sedimentation assays are as previously described [31]. Rabbit skeletal muscle actin (Cytoskeleton, AKL95) was polymerized *in vitro* as described above for pyrene-labelled actin. To assess the effect of MICAL1/PAK1 on F-actin sedimentation, recombinant human MICAL1 was incubated with or without recombinant active PAK1 in an *in vitro* phosphorylation assay as described above. 25 μl of kinase reaction was incubated with 100 μl of polymerized F-actin at room temperature and the reaction started by the addition of 100 μM NADPH. Reactions were stopped after 5 minutes by the addition of 2 mM EDTA. Reactions were transferred to polyallomer microtubes and samples centrifuged at 100,000 xG for 20 minutes at 18°C. Supernatants were carefully removed and kept on ice before the addition of 4x LDS reducing sample buffer. Pellets were resuspended in 100 μl 1x LDS reducing sample buffer and all samples boiled at 95°C for 5 minutes. Samples were resolved by SDS-PAGE and gels stained with Coomassie brilliant blue stain.

### Actin depolymerization

Actin depolymerization assays were performed using pyrene labelled actin as previously described [29, 31]. Pyrene-labelled rabbit skeletal muscle actin (Cytoskeleton, AP05), was resuspended in G-buffer (5 mM Tris pH 8.0, 0.2 mM CaCl_2_, 0.2 mM ATP) and polymerized at room temperature for 1 hr by the addition of polymerization buffer (12.5 mM KCl, 0.5 mM MgCl_2_, 0.25 mM ATP) to generate a stock of 21 μM F-actin. Prior to analysis, the F-actin stock was diluted five-fold to 0.2 mg/ml with G-buffer and rates of depolymerization analysed by monitoring fluorescence intensity at 405 nm with excitation at 360 nm using a BioTek Synergy HTX Multi-Mode Reader. To assess the effect of MICAL1/PAK1 on F-actin dynamics, recombinant human MICAL1 was incubated with or without recombinant active PAK1 in an *in vitro* phosphorylation assay as described above. 50 μl of kinase reaction was added to 200 μl of diluted F-actin, and the reaction started by the addition of 100 μM NADPH. Fluorescence intensity was measured every 30 seconds for 25 minutes.

### Proximity ligation assays

U2OS cells were transfected with FLAG-MICAL1, CDC42 G12V and PAK1^WT^ or PAK1^KD^ plasmids and left to grow overnight before fixation in 4% PFA. Detection of pS817, pS960 and total MICAL1 using a Duolink^®^ PLA kit (Millipore-Sigma) was perfomed by labelling all samples with a mouse anti-FLAG antibody (**Figure S2A**; indicated in blue) in combination with the following rabbit antibodies (**Figure S2A**; indicated in red); anti-pS817 or anti-pS960 antibodies to detect phosphorylated MICAL1, anti-FLAG antibody for total MICAL1 expression, or no antibody as a negative control for the PLA probes. Cells were stained with phalloidin and DAPI for F-actin and nuclei detection, respectively.

### Statistics

Data analysis was performed using GraphPad Prism. Statistical tests used for each data set are indicated in respective figure legends.

## Acknowledgements

This research was supported by funding to M.F.O from the Canadian Institutes of Health Research (PJT-169125), Natural Sciences and Engineering Research Council of Canada (RGPIN-2020-05388), Canada Research Chairs Program (950-231665), and the Ryerson University Faculty of Science Dean’s Research Fund. Additional funding from CRUK institutional funding to the CRUK Beatson Institute (A10419, A17196) and to M.F.O. (A18276).

---

**Figure S1.**
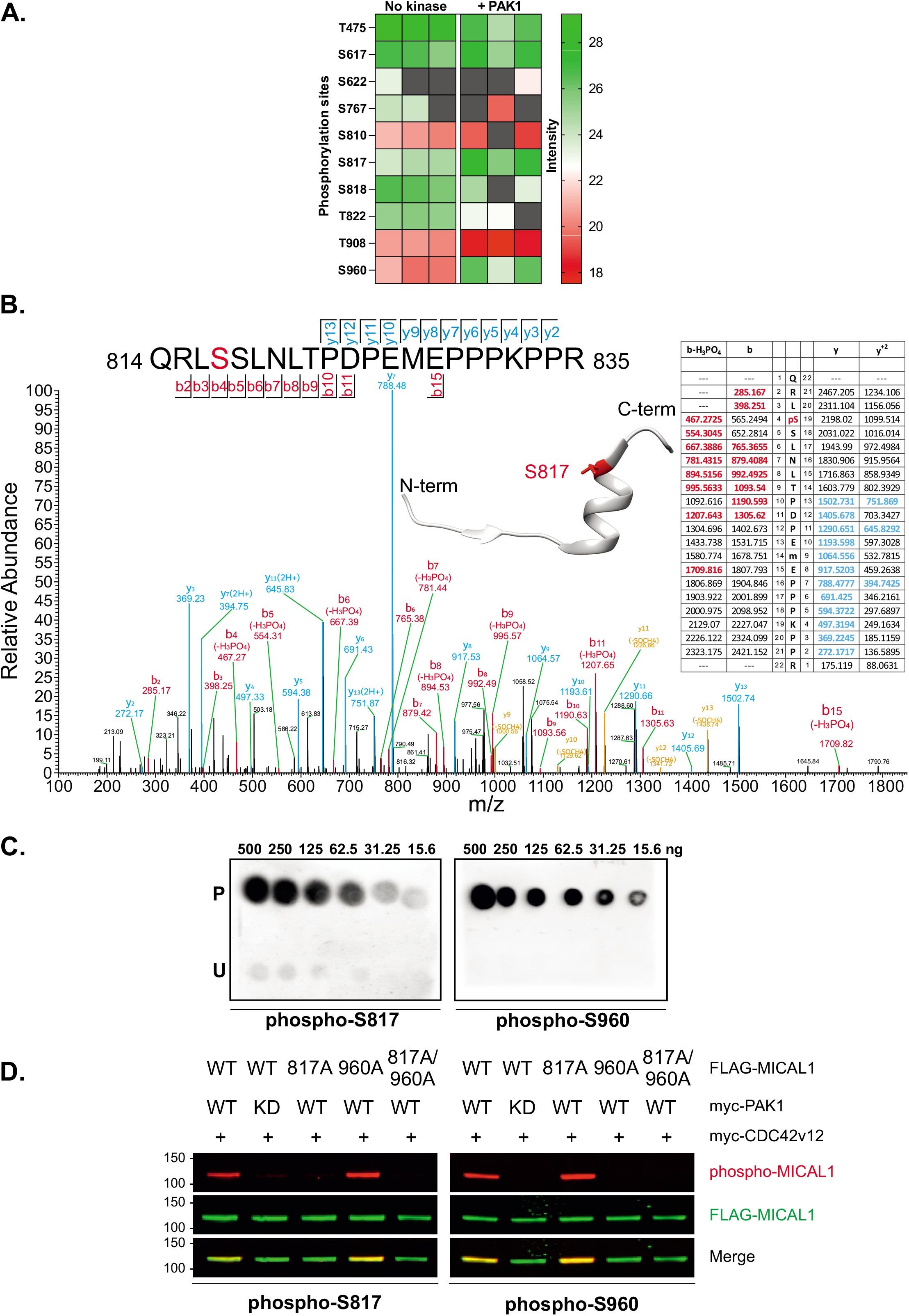
MICAL1 phosphorylation. **A.** Heat map of relative intensities of Serine and Threonine phosphorylation at indicated sites from triplicate experimental replicates for FLAG-MICAL expressed in HEK293T cells alone (no kinase) or co-expressed with CDC42 G12V and PAK1 (+ PAK1) **B.** Higher energy collision dissociation (HCD) MS/MS fragmentation spectra of MICAL1 tryptic peptide 814-835 containing S817 (red). The b and y ion series ions identified experimentally are indicated in blue and red, respectively. *Inset*, UCSF Chimera [51] modeling of the MICAL1 amino acids 796-822 indicates the S817 (red) location (Protein data base reference 6KU0) [33]. **C.** Dot blotting of phosphorylated (P) and unphosphorylated (U) peptides containing amino acid 817 (left panel) or S960 (right panel) that were spotted at the indicated quantities. Incubation of the membranes with anti-phospho-S817 (left panel) or anti-phospho-S960 (right panel) antibodies was followed by detection using a secondary rabbit-HRP antibody and chemiluminescence. **D.** HEK293T cells were transfected with indicated FLAG-MICAL1 mutants, myc CDC42 G12V and myc-tagged PAK1^WT^ or PAK1^KD^. Proteins were resolved by Western blotting and detected with anti-FLAG antibody (green), anti-phospho-S817 (left panel) or anti-phospho-S960 (right panel) MICAL1 antibodies (red).

**Figure S2.**
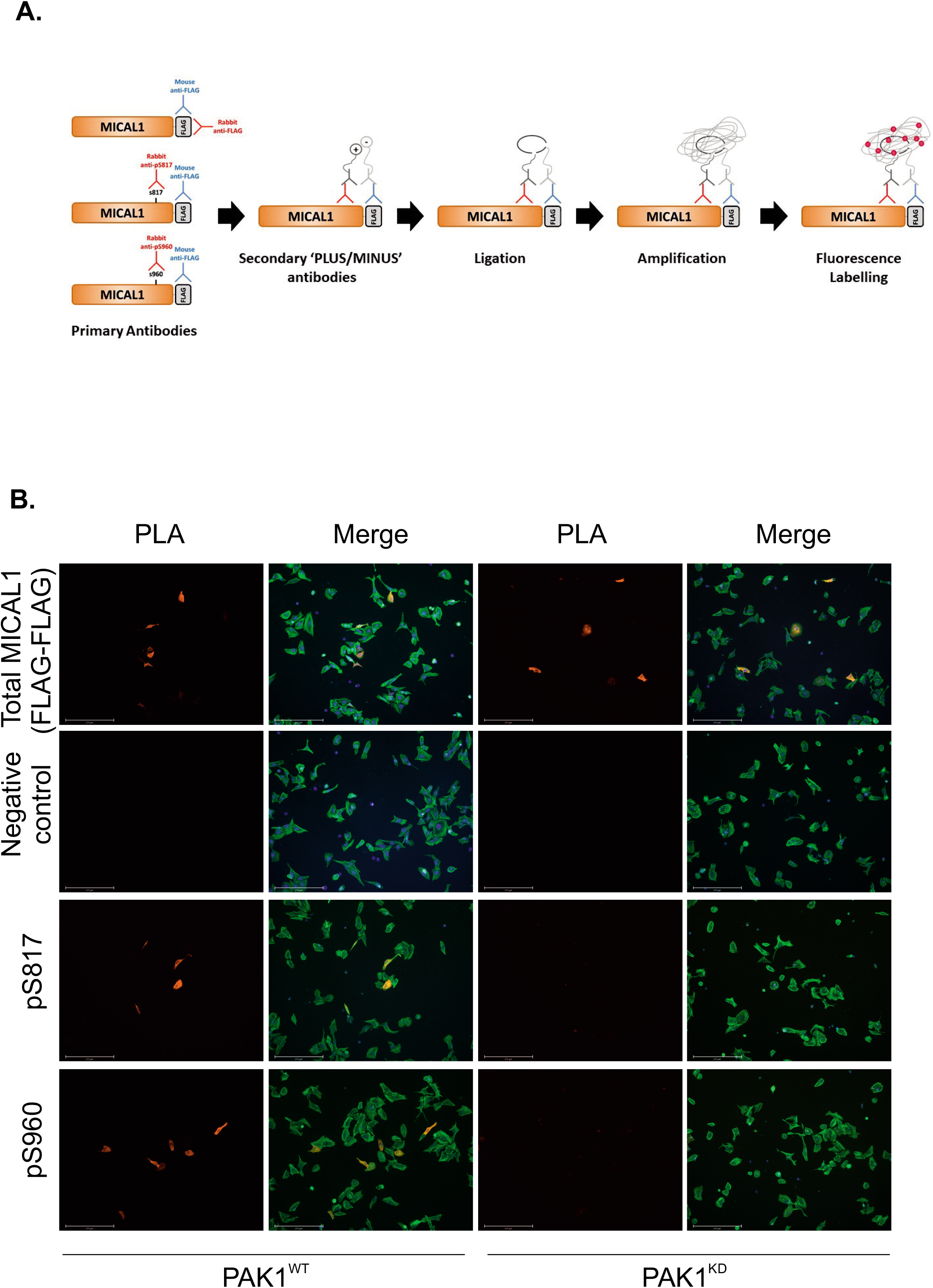
Proximity ligation assays of MICAL1 localization. **A.** Schematic diagram for proximity ligation assays (PLA) to detect total and phosphorylated MICAL1. **B.** FLAG-MICAL1 was co-expressed in U2OS cells with myc-CDC42 G12V and myc-PAK1^WT^ (left panels) or myc-PAK1^KD^ (right panels). PLA signals showing total MICAL1 (FLAG-FLAG), negative control (without rabbit anti-FLAG), phospho-S817 and S960 in red, and F-actin staining with phalloidin in green. Phosphorylated S817 and S960 were only detected in cells co-expressing myc-CDC42 G12V and myc-PAK1^WT^, consistent with the phospho-sensitivity of the antibodies. Scale bars = 275 μm.

**Figure S3.**
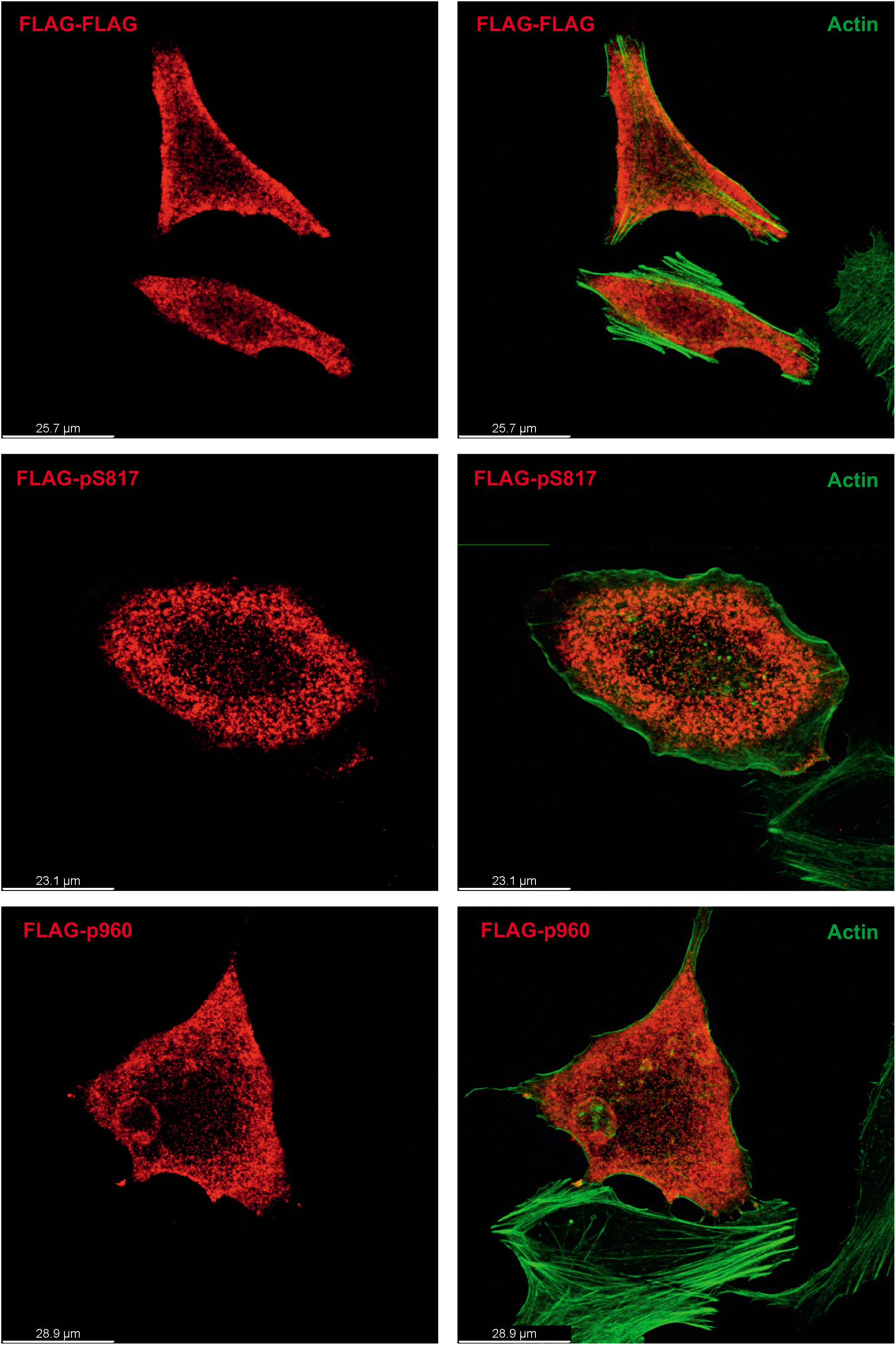
High resolution proximity ligation assays of MICAL1 localization. Maximum projection images of PLA signal (red) for total MICAL1 co-expressed with myc-CDC42 G12V and myc-PAK1^KD^ (top panels) detected with mouse plus rabbit anti-FLAG antibodies (FLAG-FLAG), or MICAL1 co-expressed with myc-CDC42 G12V and myc-PAK1^WT^ and detected with mouse anti-FLAG antibody plus rabbit anti-phospho-S817 (pS817, middle panels) or rabbit anti-phospho-S960 (pS960, bottom panels). Cells were co-stained with phalloidin (green) to detect F-actin. Scale bars indicated for each panel pair.

**Supplemental Table 1.**
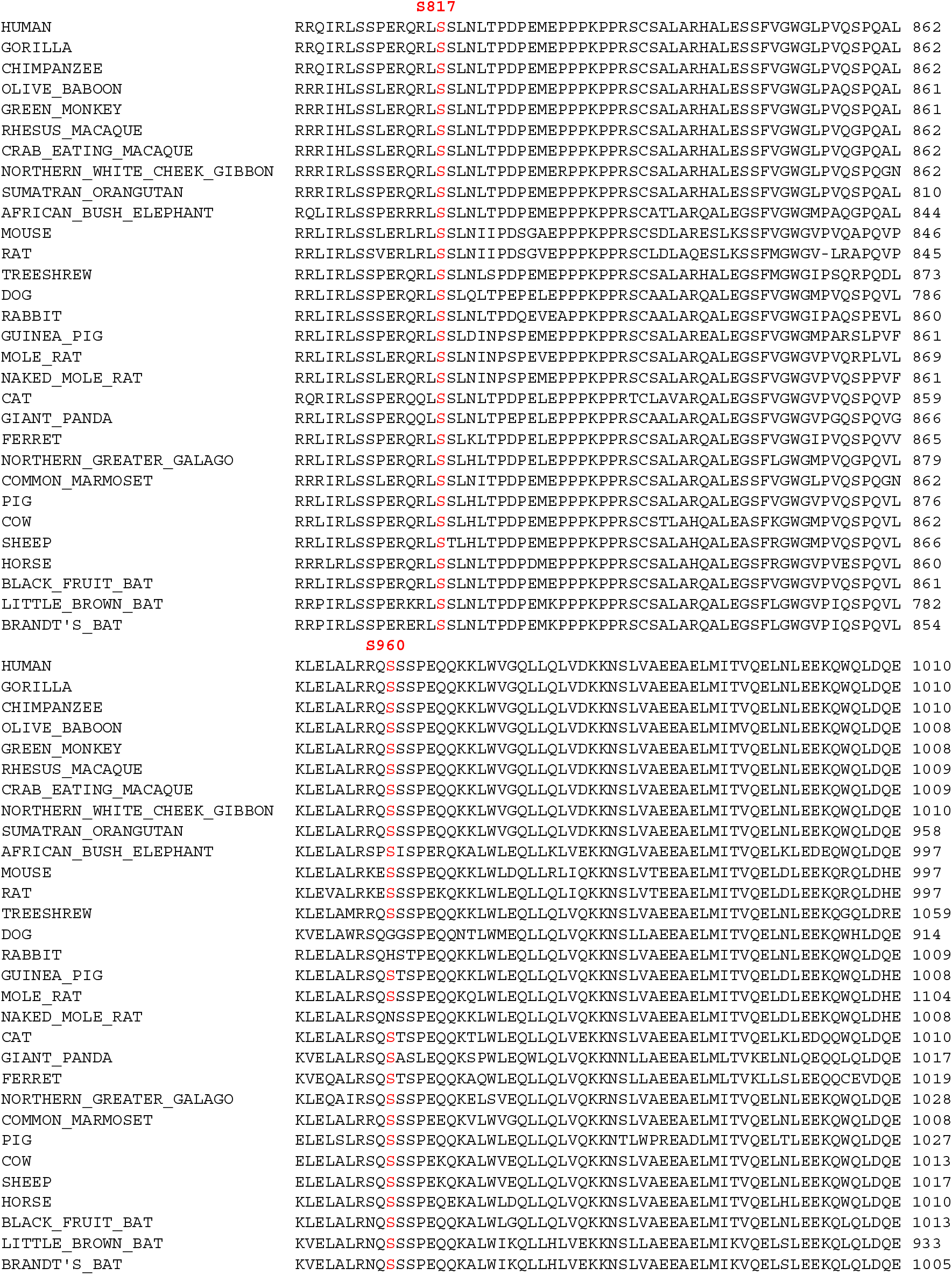
Conservation of S817 and S960 across 31 example mammalian species.

